# Substantial immune response in Omicron infected breakthrough and unvaccinated individuals against SARS-CoV-2 variants of concerns

**DOI:** 10.1101/2022.01.24.477043

**Authors:** Pragya D. Yadav, Gajanan N. Sapkal, Rima R. Sahay, Varsha A. Potdar, Gururaj R. Deshpande, Deepak Y. Patil, Dimpal A. Nyayanit, Anita M. Shete, Jayanthi Shastri, Pradeep Awate, Bharti Malhotra, Priya Abraham

## Abstract

The recent emergence of highly mutated SARS-CoV-2 Omicron variant has debilitating effect on public health system of the affected countries worldwide. Currently India is facing third wave of COVID-19 pandemic and going through a severe crisis. Within short span of time, the variant has shown high transmissibility and capability of evading the immune response generated against natural infection and vaccination. The immune escape potential of Omicron is a serious concern and further needs to be explored. In the present study, we have assessed the IgG and neutralizing antibody (NAb) response in breakthrough individuals vaccinated with two doses ChAdOx1 nCoV-19 vaccine (n=25), breakthrough individuals vaccinated with two doses of BNT162b2 mRNA vaccine (n=8) and unvaccinated individuals (n=6). All these individuals were infected with Omicron variant. The IgG antibody activity in the sera of the ChAdOx1 nCoV-19 and BNT162b2 mRNA breakthrough individuals was comparable with S1-RBD, while it was lesser in BNT162b2 mRNA breakthrough individuals with N protein and inactivated whole antigen IgG ELISA. BNT162b2 mRNA breakthrough individuals showed moderate reduction in NAb GMTs compared to ChAdOx1 nCoV-19 against Alpha, Beta and Delta. However, 3-fold higher reduction was observed with omicron variant in BNT162b2 mRNA than ChAdOx1 nCoV-19. Apparently, Alpha variant was modestly resistant to the sera of unvaccinated individuals than Beta, Delta and Omicron. Our study demonstrated substantial immune response in the individuals infected with Omicron. The neutralizing antibodies could effectively neutralize the Omicron and other VOCs including the most prevalent Delta variant.

## Background

The recent emergence of SARS-CoV-2 Variant of Concern (VOC) Omicron has caused a major spike in new COVID-19 infections worldwide [1]. Although, the Delta variant still found be the most prevalent VOC across the globe, the Omicron has displaced Delta in Southern Africa and rapidly becoming dominant in the United Kingdom and United States of America [1]. Since January 2022, India has also seen the sudden surge in COVID-19 cases with the Omicron and Delta variant. Omicron has shown higher transmissibility and immune escape as compared to the other VOCs including Alpha, Beta, and Delta leading to many reported breakthrough and re-infections across the globe. However, the severity of the disease is lesser as observed with other VOCs [2]. Further research are crucial to evaluate the immune evasion potential of the Omicron in the individuals with natural infection and/or vaccination. In this study, we have analyzed the IgG and neutralizing antibodies (NAbs) against B.1, Alpha, Beta, Delta and Omicron variants with the sera of individuals infected with the Omicron variant (B.1.1529 and BA.1).

## Materials and methods

### Study participants

The study individuals (n=39) were mainly the foreign returnees (n=28) [UAE, South/West/ East Africa, Middle east, USA and UK] and their high-risk contacts (n=11) confirmed as Omicron by next generation sequencing. The sera samples of all the individuals were collected at the post onset date of 10.5 ± 6.3 days, where oro/naso-phyrangeal swabs of eight individuals were positive for SARS-CoV-2 (Ct range: 16-29) and rest of the samples were negative. Further, the participants were grouped into three categories, breakthrough infections after two dose of vaccines [ChAdOx1 nCoV-19 (n=25); 147 ± 60 days post vaccination and BNT162b2 mRNA (n=8) 113 ± 57 days post vaccination] and unvaccinated individuals (n=6).

### Anti-SARS-CoV-2 IgG S1-RBD, N protein, Inactivated whole antigen ELISA

The samples were tested for anti-SARS-CoV-2 IgG antibodies against S1-RBD ELISA. Briefly, 96-well ELISA plates (Nunc, Germany) were coated with SARS-CoV-2 specific antigens (S1-RBD at a concentration of 1.5μg/well, N protein at a concentration of 0.5μg/well and inactivated SARS-CoV-2 antigen at a concentration of 1 μg/well in PBS pH 7.4). The plates were blocked with a Liquid Plate Sealer (CANDOR Bioscience GmbH, Germany)/ Stabilcoat (Surmodics) for two hours at 37°C. The plates were washed three times with 10 mM PBS, pH 7.4 with 0.1 per cent Tween-20 (PBST) (Sigma-Aldrich, USA). The sera of study participants (1:50 dilutions for S1-RBD and N Protein/1:100 dilutions for inactivated SARS-CoV-2 antigen) were serially four-fold diluted and added to the antigen-coated plates and incubated at 37°C for one hour. These wells were washed five times using 1× PBST and followed by addition 50 μl/well of anti-human IgG horseradish peroxidase (HRP) (Sigma) diluted in Stabilzyme Noble (Surmodics). The plates were incubated for half an hour at 37°C and then washed as described above. Further, 100 μl of TMB substrate was added and incubated for 10 min. The reaction was stopped by 1 N H2SO4, and the absorbance values were measured at 450 nm using an ELISA reader. The cut-off for the assays was set at twice of average OD value of negative control. The endpoint titer of a sample is defined as the reciprocal of the highest dilution that has a reading above the cutoff value [3].

### Plaque reduction neutralization test

Further, the plaque reduction neutralization test (PRNT50) was also performed against the B.1 (NIV2020-770, GISAID accession number: EPI_ISL_420545), Alpha [B.1.1.7, hCoV-19/India/20203522 SARS-CoV-2 (VOC) 20211012/01], Beta (B.1.351, NIV2021-893, GISAID accession number: EPI_ISL_2036294), Delta (B.1.617.2, NIV2021-1916, GISAID accession number: EPI_ISL_2400521) and Omicron (B.1.1.519, NIV2021-11828, GISAID accession number: EPI_ISL_8542931) variants which were isolated from clinical samples of COVID-19 positive cases [4]. Briefly, the plaque reduction neutralization test (PRNT) was performed in a Biosafety level 3 facility. Vero CCL-81 cell suspension (1.0 x 105 /mL/well) was added in 24-well tissue culture plates and was incubated for 16-24 hours at 37°C with 5% CO_2_ to obtain a confluent monolayer. Serum samples were inactivated at 56°C-water bath for 30 minutes. A four-fold serial dilution of serum samples were mixed with an equal volume of virus suspension containing 50-60 plaque-forming units (PFU/0.1ml) and incubated at 37°C for 1 hour. The virus–serum mixtures (0.1ml) were added onto the Vero CCL-81 cell monolayers and incubated at 37°C with 5% CO2 with intermittent shaking for uniform absorption of the virus. After 1 hour, the suspension was aspirated and the cell monolayer was gently overlaid with an overlay medium containing 2% carboxymethylcellulose (CMC) and 2X MEM containing 2% FBS (1:1). The plates were further incubated at 37°C with 5% CO2 for 4 days. The plates were terminated by decanting overlay medium followed by 0.1% saline wash and were stained with 1% amido black for 1 hour at room temperature. The plates were washed in running tap water and air-dried. Plaque number was counted and PRNT50 was defined as a reciprocal of the highest dilution of tested serum that resulted in a reduction of viral infectivity by 50% when compared to the non-neutralization control [5]. PRNT50 titre was calculated using a log probit regression analysis by SPSS (SPSS Inc., Chicago, IL).

## Results

The mean age of the ChAdOx1 nCoV-19 vaccinated individuals was 39 years [11F/14M] with thirtheen of them being asymptomatic while twelve had mild fever, sore-throat cold and cough lasting for 1-3 days. Of these, two cases were of reinfection with documented past infection in July 2020. The individuals vaccinated with two doses of BNT162b2 had mean age of 33 years [7F/1M] and only one case reported to have mild fever and sore-throat while rest seven were asymptomatic. Unvaccinated group (n=6) included all female individuals [mean age 16 years, peadiatric (n=5) and adult (n=1)]. Of these, four were asymptomatic while two peadiatric cases were mildly symptomatic.

The geometric mean titres (GMTs) of the S1-RBD IgG antibodies in the sera of the ChAdOx1 nCoV-19 [1179] and BNT162b2 mRNA [1383] breakthrough individuals showed no significant difference. However, 2.5 to 3.6 times reduction in the GMTs was observed with N protein and inactivated whole antigen IgG ELISA in the sera of BNT162b2 mRNA breakthrough individuals compared to the ChAdOx1 nCoV-19. In the unvaccinated group, varied response in the GMTs of IgG antibodies were observed with S1-RBD (166.9), N protein (30) and inactivated whole antigen (357.8) [Figure-1 A, B, C].

**Figure-1:**
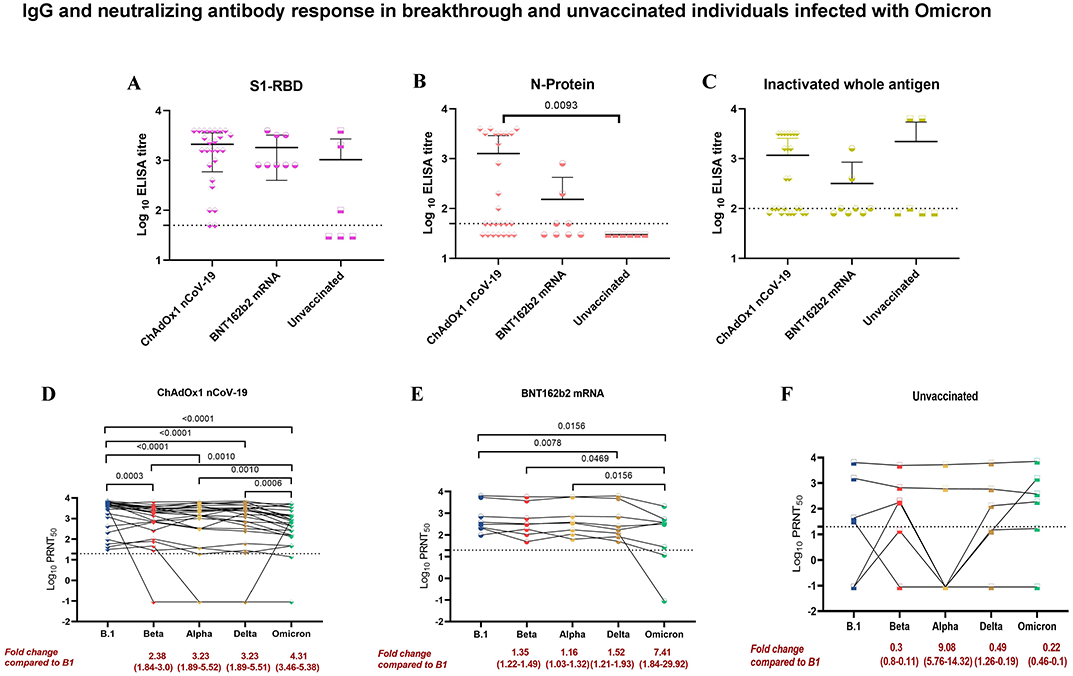
IgG antibody response in breakthrough and unvaccinated individuals was assessed using ELISA A) S1-RBD B) N protein C) Inactivated whole antigen. The IgG antibody titers of the different groups were compared using the two-tailed Kruskal Wallis test and the p-value less than 0.05 were considered to be statistically significant. The neutralizing antibody response among D) ChAdOx1 nCoV-19 breakthrough E) BNT162b2 mRNA breakthrough and F) unvaccinated individuals against B.1, Alpha, Beta, Delta and Omicron were determined using the plaque reduction neutralization test (PRNT50). The titers amongst the groups were compared with B.1 using the Wilcoxon matched-pairs signed-rank test to assess the statistical significance.

The GMTs of neutralizing antibodies of ChAdOx1 nCoV-19 breakthrough individuals showed significant fold-reductions compared to B.1 against Alpha (3.23), Beta (2.38), Delta (3.23) and Omicron (4.31) variants respectively. Similarly, BNT162b2 mRNA breakthrough individuals demonstrated significant fold-reduction in GMTs of 1.52 and 7.41 for Delta and Omicron respectively. While, non-significant fold-reduction was observed with Alpha (1.16) and Beta (1.35). In contrary, Alpha variant (9.08) was modestly more resistant to neutralization than Beta (0.3), Delta (0.49) and Omciron (0.22) in the unvaccinated individuals compared to B.1 [Figure-1 D,E, F].

## Discussion

The main limitation of this study are lesser participants in the unvaccinated group and the shorter window period post infection. This could be the important reason for the low immune response specifically in the unvaccinated individuals against Omicron.

Our study suggest a 3-fold reduction in the NAb titres in BNT162b2 mRNA breakthrough individuals as compared with ChAdOx1 nCoV-19.The earlier studies from other groups demonstrated that immune response generated through natural infection or vaccination showed weaker immune response against Omicron [6–10]. Our study demonstrated that the individuals infected with Omicron have significant immune response which could neutralize not only the Omicron but also the other VOCs including most prevalent Delta variant. This suggest that the immune response induced by the Omicron could effectively neutralize the Delta variant making the re-infection with Delta less likely, thereby displacing the Delta as dominant strain. This emphasizes the need for the omicron specific vaccine strategy.

## Ethical approval

The study was approved by the Institutional Biosafety Committee and Institutional Human Ethics Committee of ICMR-NIV, Pune, India under the project titled “Assessment of immunological responses in breakthrough cases of SARS-CoV-2 in post COVID-19 vaccinated group” [No.NIV/IEC/May/2021/D-10 dated May 20, 2021].

## Author Contributions

PDY contributed to study design, data analysis, interpretation and writing and critical review. GNS, RRS, VAP, GRD, DAN, DYP, AMS, PA, JS, BM contributed to data collection, interpretation, writing and critical review. PDY, RRS, DYP and PA contributed to the critical review and finalization of the paper.

## Acknowledgement

Authors are thankful to the support provided by Dr. Kiran Bhise, In-charge COVID care Centre Pune; Dr. Balaji Barure, District Surveillance officer (DSO), Latur; Dr. Bodke, DSO Osmanabad; Dr. Sachin Patil, DSO Satara for the coordination of the sample collection and shipment. The authors would like to gratefully acknowledge the staff of ICMR-NIV, Pune including Mrs. Asha Salunkhe, Mr. Chetan Patil Dr. Rajlaxmi Jain, Mr. Prasad Sarkale, Mr. Hitesh Dighe, Mr. Annasaheb Suryavanshi, Mr. Manjunath Holeppanawar, Mr. Sanjay Thorat, Mrs. Priyanka Waghmare, Mrs. Poonam Bodke, Mrs. Shilpa Ray for extending the excellent technical support. The authors also acknowledge the contribution from the team of National Influenza Centre (NIC), ICMR-NIV, Pune.

## References

1. World Health Organization. WHO coronavirus (COVID-19) dashboard. https://covid19.who.int/ Accessed 18 January 2022.

2. Wolter N, Jassat W, Walaza S, et al. Early assessment of the clinical severity of the SARS-CoV-2 Omicron variant in South Africa. medRxiv. 2021 Jan 1.

3. Ella R, Reddy S, Jogdand H, et al. Safety and immunogenicity of an inactivated SARS-CoV-2 vaccine, BBV152: interim results from a double-blind, randomised, multicentre, phase 2 trial, and 3-month follow-up of a double-blind, randomised phase 1 trial. Lancet Infect Dis. 2021 Mar 8.

4. Yadav PD, Gupta N, Potdar V, et al. An in vitro and in vivo approach for the isolation of Omicron variant from human clinical specimens. bioRxiv. 2022 Jan 1.

5. Deshpande GR, Sapkal GN, Tilekar BN, et al. Neutralizing antibody responses to SARS-CoV-2 in COVID-19 patients. Indian J Med Res. 2020;152(1-2):82.

6. Cele S, Jackson L, Khoury DS, et al. SARS-CoV-2 Omicron has extensive but incomplete escape of Pfizer BNT162b2 elicited neutralization and requires ACE2 for infection. medRxiv. 2021.

7. Andrews N, Stowe J, Kirsebom F, et al. Effectiveness of COVID-19 vaccines against the Omicron (B.1.1.529) variant of concern. medRxiv, 2021.

8. Wilhelm A, Widera M, Grikscheit K, et al. Reduced Neutralization of SARS-CoV-2 Omicron Variant by Vaccine Sera and Monoclonal Antibodies. medRxiv, 2021.

9. Roessler A, Riepler L, Bante D, et al. SARS-CoV-2 B.1.1.529 variant (Omicron) evades neutralization by sera from vaccinated and convalescent individuals. medRxiv, 2021.

10. Carreno JM, Alshammary H, Tcheou J, et al. Activity of convalescent and vaccine serum against SARS-CoV-2 Omicron. Nature. 2021:1–8.

